# TAMPA: interpretable analysis and visualization of metagenomics-based taxon abundance profiles

**DOI:** 10.1101/2022.04.28.489926

**Authors:** Varuni Sarwal, Jaqueline Brito, Serghei Mangul, David Koslicki

## Abstract

Metagenomic taxonomic profiling aims to predict the identity and relative abundance of taxa in a given whole genome sequencing metagenomic sample. A recent surge in computational methods that aim to accurately estimate taxonomic profiles, called taxonomic profilers, have motivated community driven efforts to create standardized benchmarking datasets and platforms, standardized taxonomic profile formats, as well as a benchmarking platform to assess tool performance. While this standardization is essential, there is currently a lack of tools to visualize the standardized output of the many existing taxonomic profilers, and benchmarking studies rely on a single value metrics to compare performance of tools and compare to benchmarking datasets. Here we report the development of TAMPA (**Ta**xonomic **m**etagenome **p**rofiling evalu**a**tion), a robust and easy-to-use method that allows scientists to easily interpret and interact with taxonomic profiles produced by the many different taxonomic profiler methods beyond the standard metrics used by the scientific community. We demonstrate the unique ability of TAMPA to provide important biological insights into the taxonomic differences between samples otherwise missed by commonly utilized metrics. Additionally, we show that TAMPA can help visualize the output of taxonomic profilers, enabling biologists to effectively choose the most appropriate profiling method to use on their metagenomics data. TAMPA is available on GitHub, Bioconda and Galaxy Toolshed at https://github.com/dkoslicki/TAMPA, and is released under the MIT license.

## Introduction

Metagenomics has become an essential tool to study microbiomes due to improvements in technology and bioinformatic algorithms. Taxonomic metagenome profiling aims to predict the identity and relative abundances of taxa in a given whole genome sequencing (WGS) metagenomic sample. A recent surge in computational methods that aim to accomplish this, called taxonomic profilers, have motivated community-driven efforts to create standardized benchmarking datasets ^1–3^, standardized taxonomic profile formats ^4^, as well as a benchmarking platform to assess tool performance on simulated data ^5^. While this standardization is essential, there is currently a lack of tools to visualize the standardized output of the many existing taxonomic profilers, and benchmarking studies rely on a single value metrics to compare performance of tools and compare to benchmarking datasets. Indeed, the only two such WGS taxonomic profiling visualization and analysis tools that do exist are either integrated into a single taxonomic profiling method ^6^, or else lack the flexibility and interpretability for the analysis and visualization of multiple taxonomic profiles ^7^. Neither of these methods are designed for or compatible with the community-driven output formats previously mentioned.

Despite the availability of flexible and interactive visualization tools in the area of amplicon microbial analysis (such as 16S rRNA studies), similar methods are yet to be developed for WGS metagenomics. For example, metacoder ^8^ is a tool that allows for visualizing, analyzing, and manipulating amplicon microbial data. However, metacoder is not designed for WGS metagenomic analyses and cannot be used for analysis and visualizing metagenomic taxonomic profiles due to amplicon analyses relying on Operational Taxonomic Units, a concept that is not relevant to metagenomic studies. Similarly, the recently published preprint for the software package EMPress^9^ is an interactive phylogenetic tree viewer not explicitly intended for the visualization of WGS taxonomic profiles.

Additionally, lack of tools that provide an interpretable visualization of multiple taxonomic profiles limits the ability of the biomedical community to select a tool. As such, when WGS metagenomic data is generated and a scientist wishes to determine which of the dozens ^10–23^ of taxonomic profilers to use, they typically rely on benchmark studies ^1,24,25^. These benchmark studies often use simulated data that does not accurately reflect their samples of interest. Alternatively, they can run their own simulation and benchmarking study tailor to their use-case, but this requires significant time investment ^2^. Often scientists resort to simply picking a familiar tool regardless of its performance characteristics. Given the substantial variability in the performance of taxonomic profiling tools ^1,24,25^, this may result in misinterpretation of their data and can potentially lead to unfortunate situations where utilizing a single taxonomic profiling tool can lead to an interpretation of data ^26^ (i.e. presence of Bubonic plague in the New York subway system) that is later to be found to be inaccurate ^27^.

Furthermore, after a taxonomic profiler is utilized, scientists can encounter difficulty when interpreting statistical measures of differences between the estimated taxonomic frequencies and the ground truth, as well as when comparing differences between tools. This difficulty is further compounded when new or emerging statistical measures are used to characterize differences between metagenomic samples, where confusion can arise about how to interpret such measures ^9, 28^.

To empower biomedical researchers with a robust and easy to use metagenomic taxonomic profile analysis and visualization platform, we have developed a software package TAMPA (Taxonomic metagenome profiling evaluation). Our platform assists scientists in contextualizing, assessing, and extracting insight from taxonomic profiles produced by multiple taxonomic profilers when applied to either real or simulated data. TAMPA is designed to allow users to effectively analyze one or more taxonomic profiles produced by any of the numerous taxonomic profiling methods. Additionally, TAMPA can operate on the widely utilized and community developed BIOM ^29^ and CAMI ^1^ profiling formats. We demonstrate the utility of TAMPA by showing how it illuminates the important biological differences between samples and conditions otherwise missed by commonly utilized statistical metrics. When gold standard taxonomic profiles are available, we show how TAMPA can augment existing benchmarking platforms such as OPAL by being incorporated within the tool and providing an interpretable visualization of the profiles ^5^. Additionally, we show that TAMPA can enable biologists to choose an appropriate profiling method to use on their real data when a ground truth taxonomic profile is not available, since TAMPA allows users to quickly ascertain similarities or differences in predictions made by multiple taxonomic tools.

## Results

### TAMPA: interpretable analysis and visualization of metagenomics-based taxon abundance profiles

TAMPA is a computational tool that allows the user to effectively visualize one or more taxonomic profiles produced by taxonomic profiling methods. TAMPA contextualizes, assesses, and extracts insight from multiple taxonomic profiler results. The taxonomic profile files are provided to TAMPA as an input and it produces a graph with the relative abundances of taxa in a given sample. The input profile files are first parsed and converted into objects. Then, the ete toolkit is used to convert the python objects into trees. We have added several command line options to improve the visualization of the tree. TAMPA allows for users to choose among multiple graph layout formats, including pie, bar, circle and rectangular. Users can further customize the graph by choosing the scaling options for the graph (log, sqrt, power), and other parameters such as the branch vertical margin, leaf separation, label font size, figure width and height, and image resolution. In the case that the number of samples is very large and the graph becomes crowded, users can choose to display only the nodes with abundance higher than a particular threshold, and/or add labels to specific parts of the graph such as only the leaf nodes. Users can also choose if they want to plot the L1 error or normalize the relative abundances of the samples. TAMPA allows users to analyze one or more samples of interest, and allows for the analysis of both single input taxonomic profiles, as well as input profiles with a ground truth. Users can also choose to decide alternate taxonomies, and restrict visualization to a particular taxonomic rank. It can be used to study the impact of filtering low abundance taxa. While the default database used for reading the input is the ncbi taxdump database, the users can specify a different database dump file. TAMPA is run using a command line interface, and takes the profile files as an input and provides a plot with the relative abundances of the taxa as the output.

To demonstrate the ability of TAMPA to provide an interpretable analysis and visualization of metagenomics-based taxon abundance profiles, we apply it to the results of three profiles from the publicly available CAMI dataset^1–3^: MetaPhyler ^14^; mOTU ^15^; and Taxy-Pro ^30^. We demonstrate three major ways in which TAMPA provides a novel way to visualize the outputs of existing profilers and visualization platforms.

### TAMPA enables effective comparison of the outputs of multiple profilers

First, TAMPA can be used to compare the outputs of multiple profilers and reveal insight even when traditional metrics report no differences. TAMPA does this by identifying which specific clades contributed to metric values, thus revealing biological differences that could otherwise be overlooked when looking only at metric values. We choose two profilers with an identical UniFrac score on a particular sample, Taxy-Pro and Metaphyler, and demonstrate the specific differences in their predictions of taxonomic profiles using TAMPA on the phylum level (Figure 1a), as well as other taxonomic levels (Figure S1-S5).

**Figure 1:**
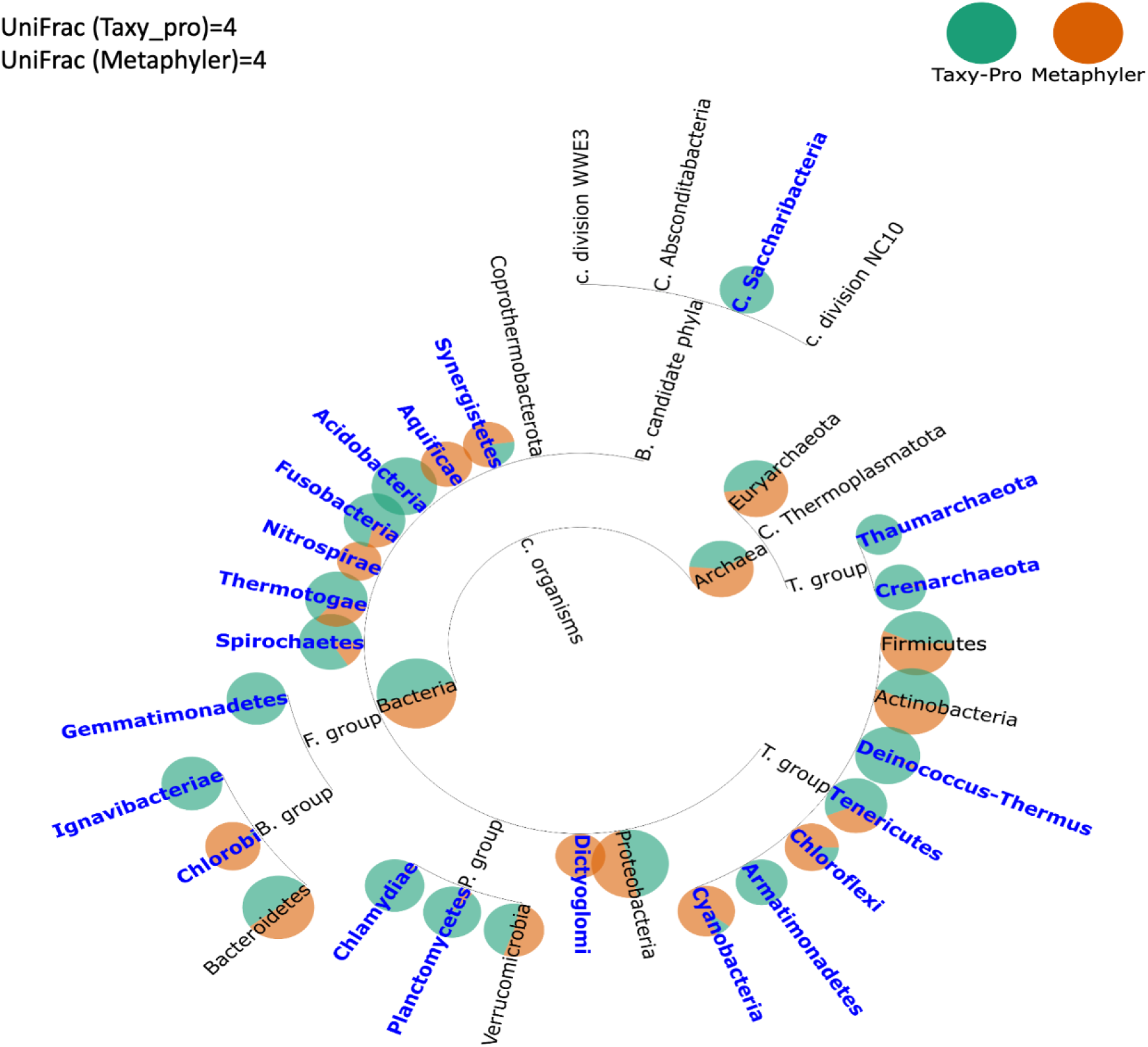
Visualization of the taxonomic profiles of **tools with identical UniFrac scores** of 4, Taxy_pro vs Metaphyler using TAMPA on the CAMI dataset at the phylum level. The size of discs represents the total amount of relative abundance at the corresponding clade in the ground truth, or the tool prediction if that clade is not in the ground truth. If the tool predictions agree, a disc is colored half orange and half teal. The proportion of teal to orange changes with respect to the disagreement in prediction of that clade’s relative abundance between the two tools being compared. Highlighted text represents clades where the difference between the relative abundances of the prediction and ground truth exceeds 30%.

**Figure 2:**
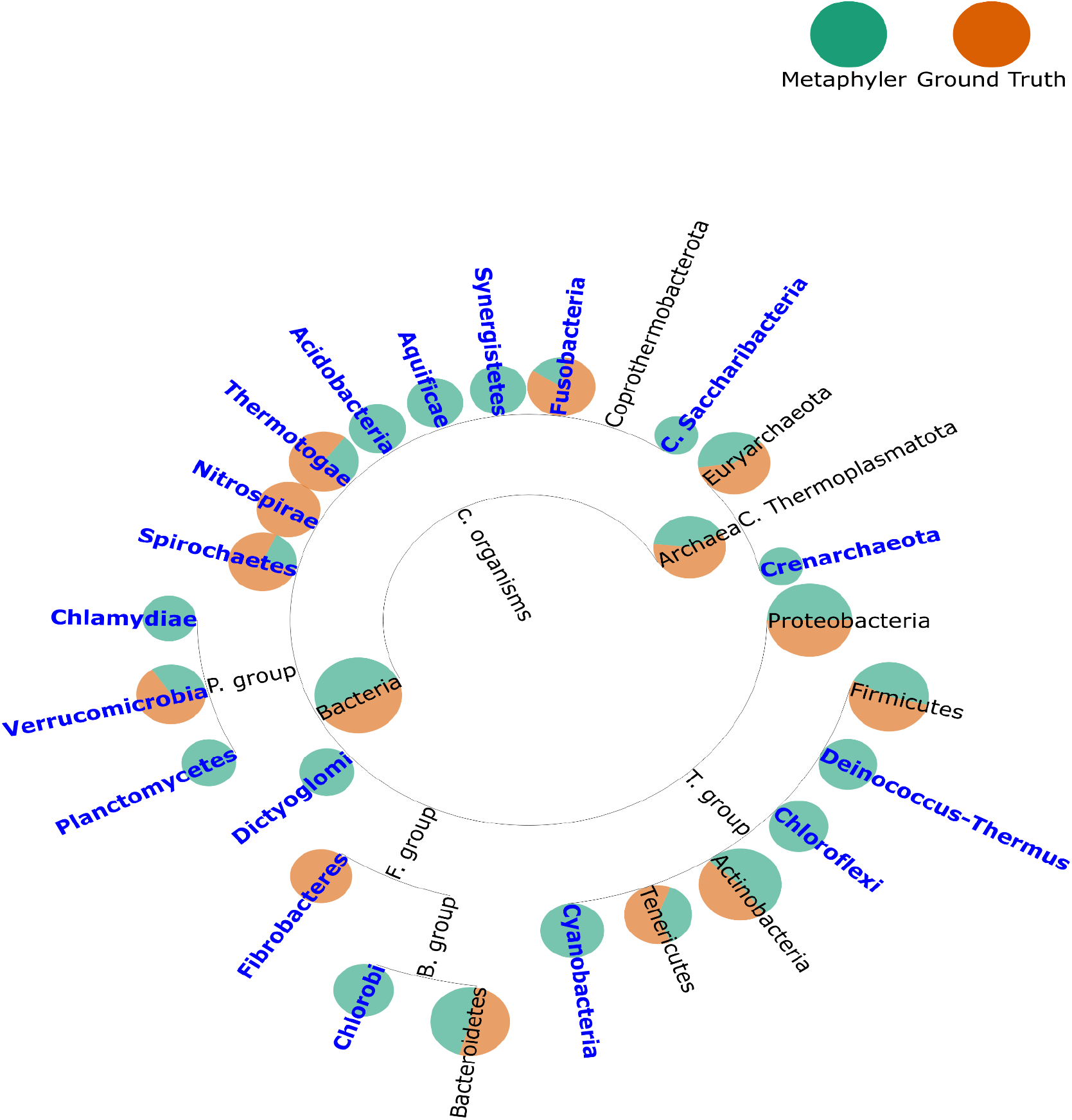
Visualization of the taxonomic profile of a **top performing CAMI tool** in terms of L1 norm, Metaphyler vs the ground truth using TAMPA on the CAMI dataset at the phylum level. The size of discs represents the total amount of relative abundance at the corresponding clade in the ground truth, or the tool prediction if that clade is not in the ground truth. If the tool predictions agree, a disc is colored half orange and half teal. The proportion of teal to orange changes with respect to the disagreement in prediction of that clade’s relative abundance between the two tools being compared. Highlighted text represents clades where the difference between the relative abundances of the prediction and ground truth exceeds 30%.

**Figure 3:**
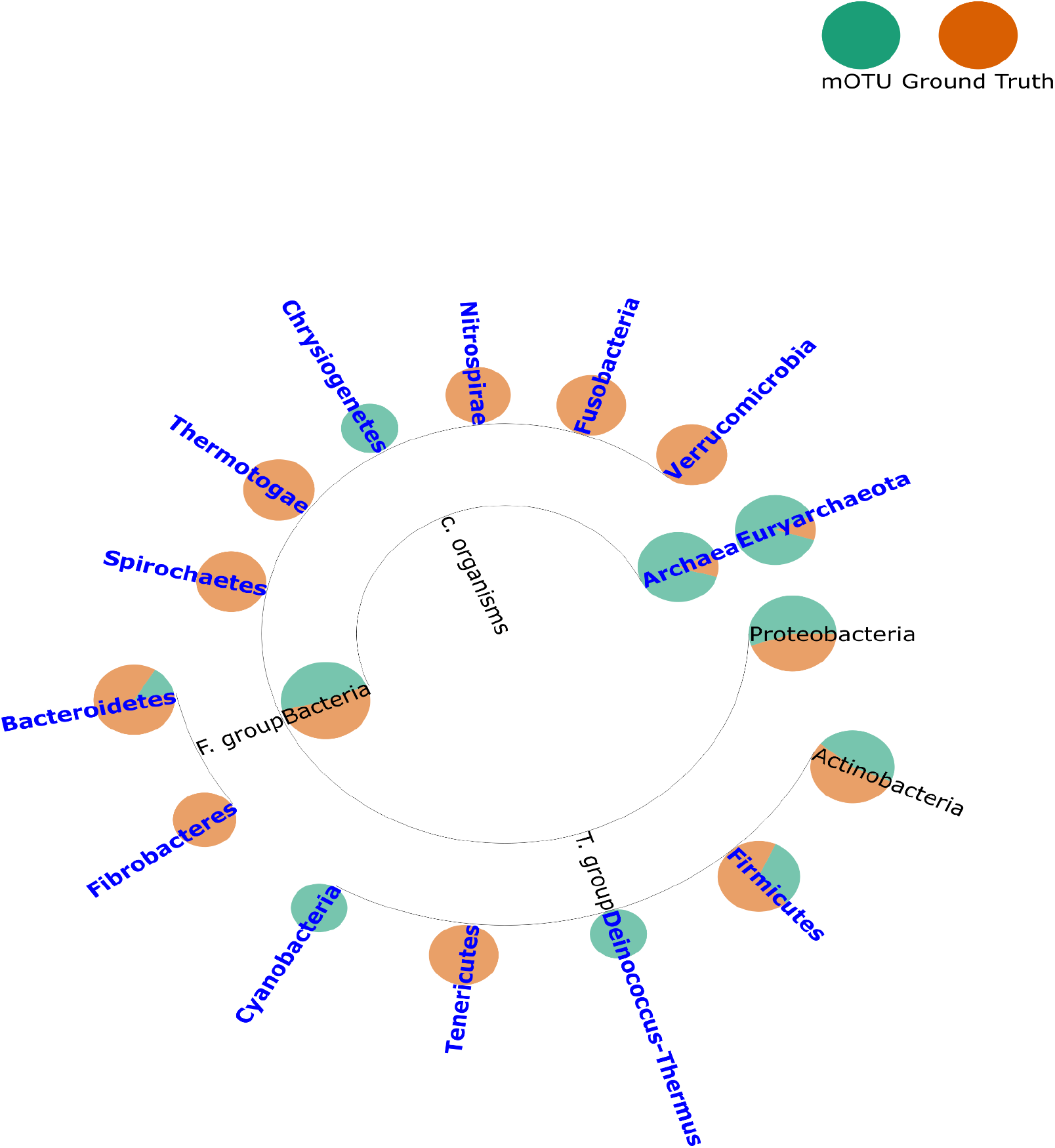
Visualization of the taxonomic profile of the **lowest performing tool in terms** of L1 norm, mOTU vs the ground truth using TAMPA on the CAMI dataset at the phylum level. The size of discs represents the total amount of relative abundance at the corresponding clade in the ground truth, or the tool prediction if that clade is not in the ground truth. If the tool predictions agree, a disc is colored half orange and half teal. The proportion of teal to orange changes with respect to the disagreement in prediction of that clade’s relative abundance between the two tools being compared. Highlighted text represents clades where the difference between the relative abundances of the prediction and ground truth exceeds 30%.

Second, even when tools performance is distinguishable by traditional numerical metrics, TAMPA can be used to quickly ascertain how tool predictions differ from the ground truth profile. For example, we chose both the top (Figure 1b) and bottom (Figure 1c) performing tools in terms of the L1 Norm, according to the CAMI challenge: Metaphyler, and mOTU, and demonstrate that TAMPA can illuminate important biological differences between the two tools and the ground truth at the phylum level (Figure 1b,c), as well as at all other taxonomic ranks (Figure S6-S15).

### TAMPA augments existing benchmarking platforms

TAMPA can be used to augment existing benchmarking platforms. We have integrated TAMPA into the taxonomic profiling benchmarking platform OPAL (^5^) in order to provide biological insight when scientists and tool developers aim to benchmark and compare taxonomic profilers (Figure S16). OPAL is a popular web-based tool used to compute commonly used performance metrics for profiler outputs. While OPAL provides global metrics and visualizations, it is unable to provide specific information on the taxonomic differences in the profiles. With the inclusion of TAMPA in OPAL, users can now quickly ascertain the performance of the tools being analyzed at a level of resolution not possible before. For example, by utilizing the figures returned by TAMPA, a user can quantify tool performance on a particular taxonomic clade of interest. Based on our results (eg., Supplementary Figure S1), we show that TAMPA can highlight important taxonomic differences easily missed by statistical metrics, thus enabling biologists to choose the most appropriate profiling method to use on their data.

## Discussion

Metagenomics has emerged as a technology of choice for analyzing microbial communities, with thousands of WGS metagenomic samples being produced annually ^31^. Taxonomic profiling is an important first step in analyzing these kinds of data. Hence, TAMPA will be of broad interest to all scientists engaged in such research as this important first step will now be easily interpreted, thus allowing scientists to quickly contextualize, assess, and extract insight from taxonomic profiles instead of relying primarily on statistical summaries or manual manipulation. Indeed, TAMPA was effectively applied in the second round of the Critical Assessment of Metagenome Interpretation (CAMI) competition^32^ where it was used to visualize the most difficult to correctly classify taxa. We present this tool as an example of how adoption of the standards put forward by the CAMI consortium can help facilitate tool development.

## Code availability

TAMPA is provided in a platform independent fashion via Bioconda ^33^: Bioconda link: https://anaconda.org/vsarwal/tampa as well as integrated into the Galaxy Toolshed ^34^ for easy “point and click” analysis for less computationally inclined users: Galaxy Toolshed link: https://toolshed.g2.bx.psu.edu/repository?repository_id=7b5054a8c1e84051

The source code is available at: https://github.com/dkoslicki/TAMPA All code required to produce the figures and analysis performed in this paper are freely available at: https://github.com/Addicted-to-coding/TAMPA_publication

## Methods

TAMPA was run on the profiling datasets generated in the CAMI challenge. The profile files were extracted from the github repo of the CAMI challenge: https://github.com/CAMI-challenge/firstchallenge_evaluation/tree/master/profiling/data/profile_submissions. The description.property file, found in the corresponding subdirectory of each tool athttps://github.com/CAMI-challenge/firstchallenge_evaluation/tree/master/profiling/data/profile_submissions was used to map the anonymous name to the tool name. We limited our analysis to Sample 1 of the high complexity dataset, denoted by CAMI_HIGH_S001. We studied tools with the highest and lowest precision, recall and UniFrac score. The following command was used to run TAMPA: python src/profile_to_plot.py -i tool.profile -g ground_truth rank -b basename -k linear -o.

TAMPA inputs several command line options from the user, such as the input and ground truth profile, output file name and sample interest, and visualization options including fontsize, labelsize and label width, layout and leaf separation. It then parses them into parameters for the plot. Depending on the user input for the different command line parameters, the ete3 toolkit is used to generate a tree which is saved after the tool finishes running.

### Data availability

TAMPA was run on the .profile files produced by the top and bottom performing taxonomic profilers. The taxonomic profiles represent the taxonomic identities and relative abundances of microbial community members from metagenome samples. The profiling files used to run TAMPA are freely available on the github repo of the CAMI challenge: https://github.com/CAMI-challenge/firstchallenge_evaluation/tree/master/profiling/data/profile_submissions.

## Supplementary Materials

### Supplementary Figures

**Figure S1:**
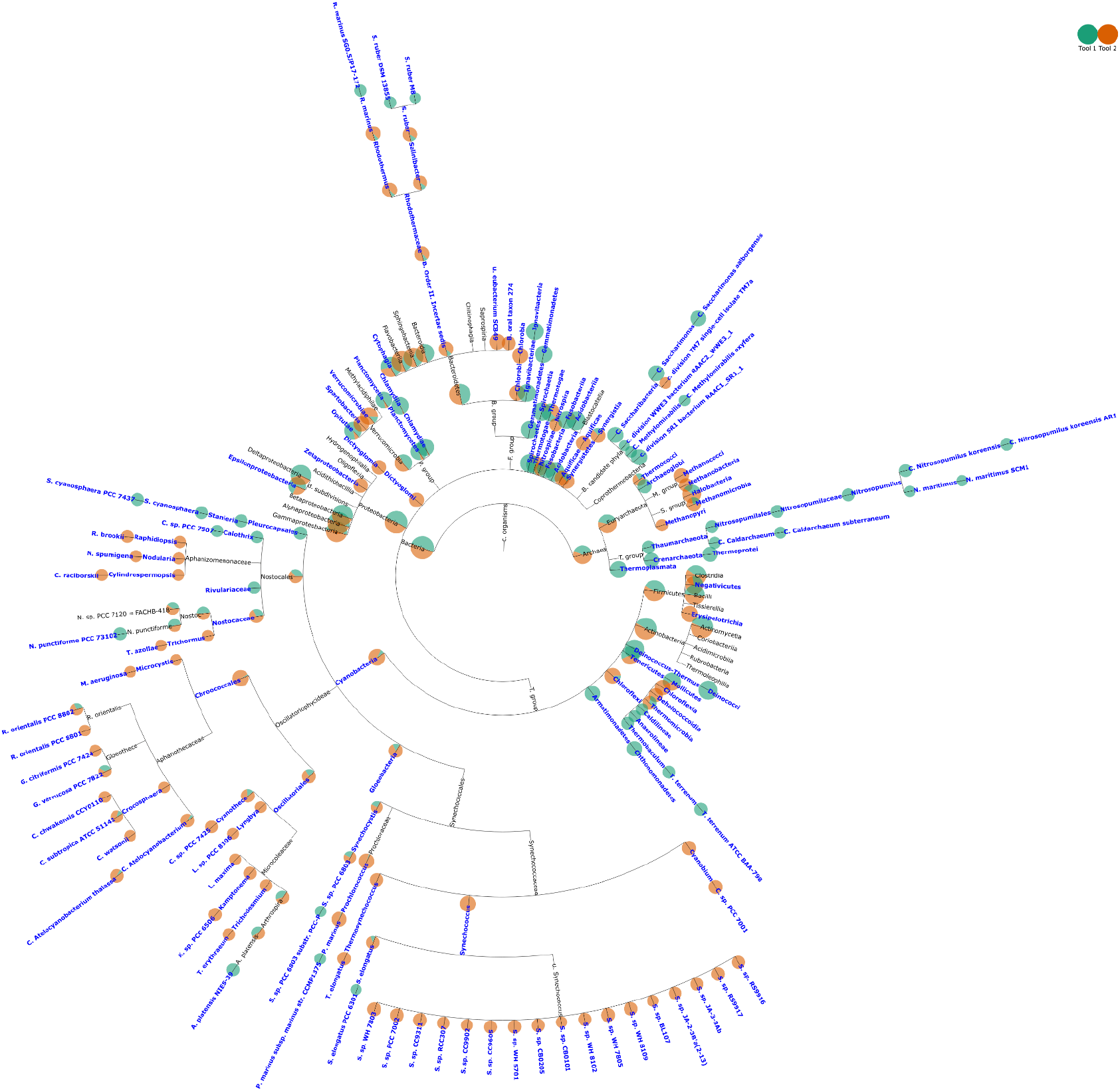
Visualization of the taxonomic profiles of tools with identical UniFrac scores of 4, Taxy_pro vs Metaphyler using TAMPA on the CAMI dataset at the class rank. Note the differences in taxa predictions even though the tools have identical UniFrac scores.

**Figure S2:**
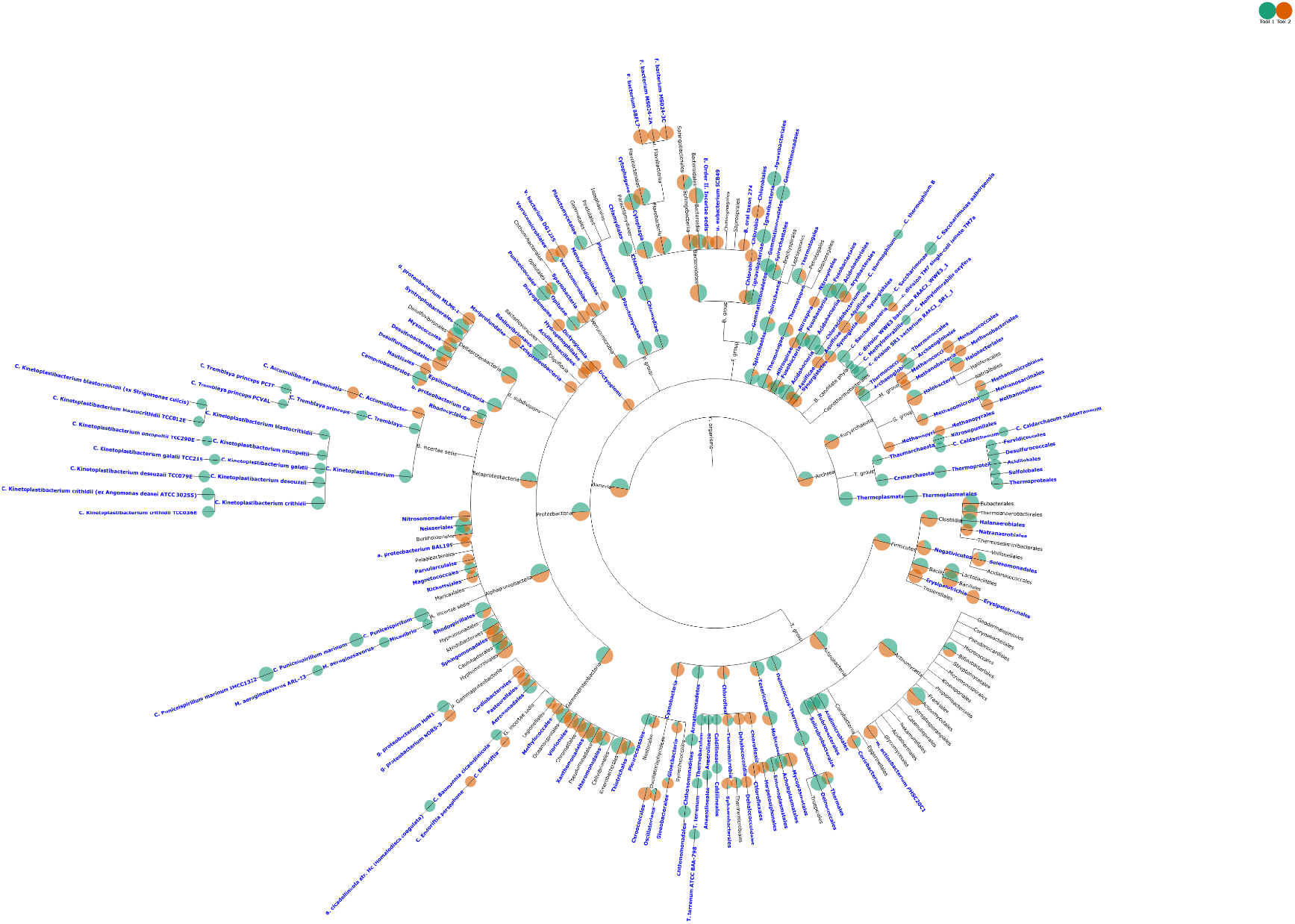
Visualization of the taxonomic profiles of tools with identical UniFrac scores of 4, Taxy_pro vs Metaphyler using TAMPA on the CAMI dataset at the order rank.

**Figure S3:**
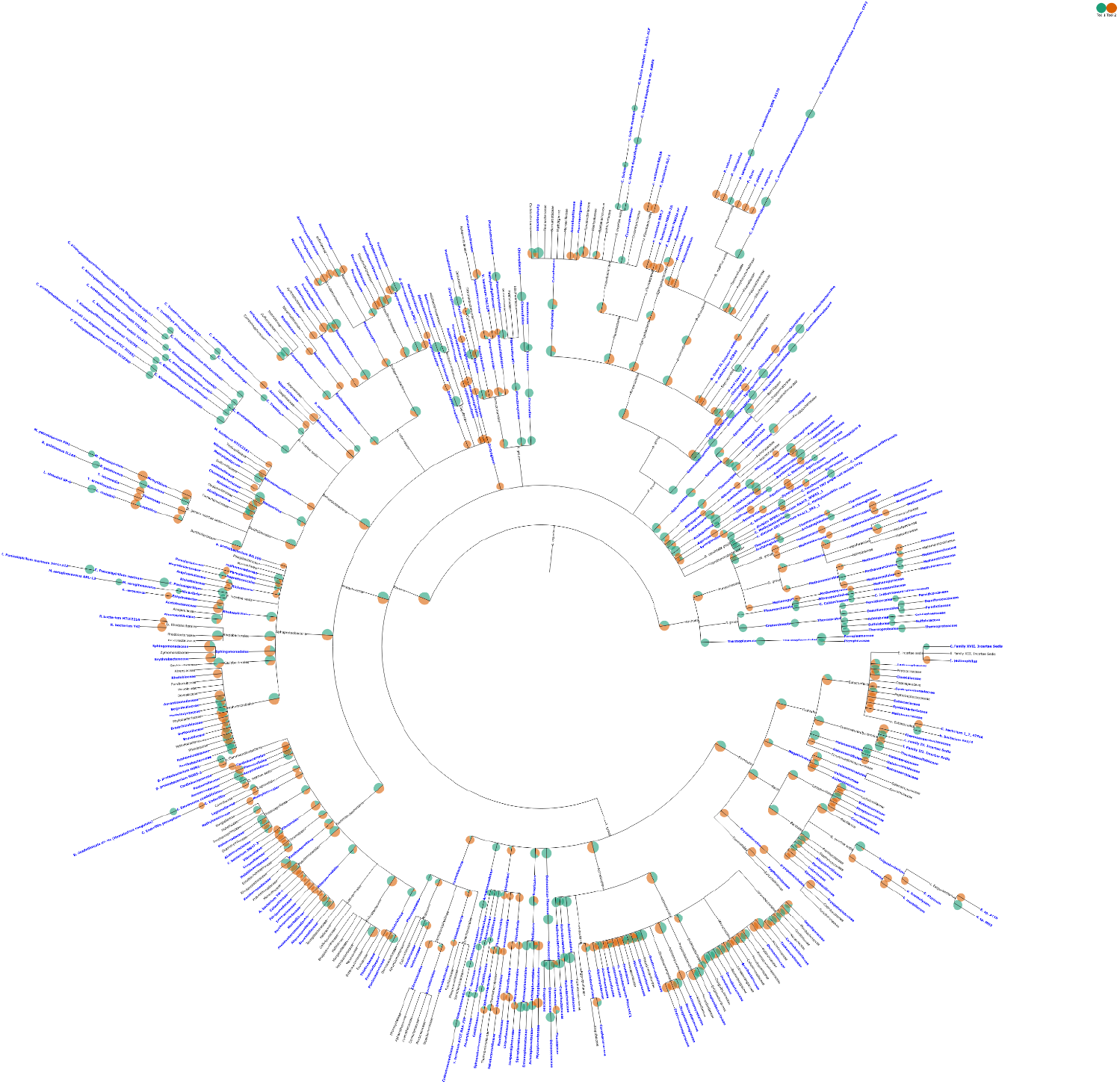
Visualization of the taxonomic profiles of tools with identical UniFrac scores of 4, Taxy_pro vs Metaphyler using TAMPA on the CAMI dataset at the family rank

**Figure S4:**
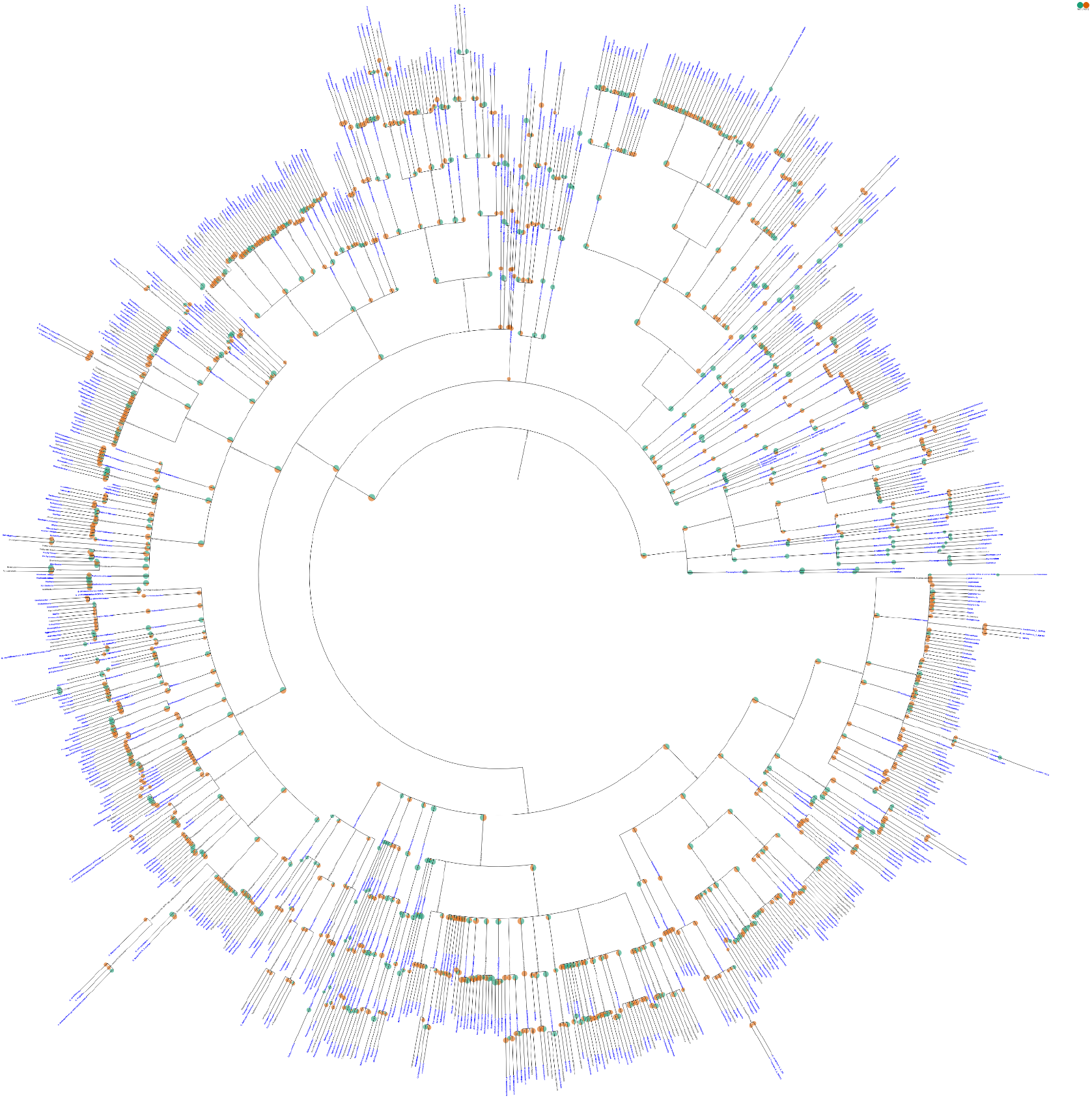
Visualization of the taxonomic profiles of tools with identical UniFrac scores of 4, Taxy_pro vs Metaphyler using TAMPA on the CAMI dataset at the genus rank

**Figure S5:**
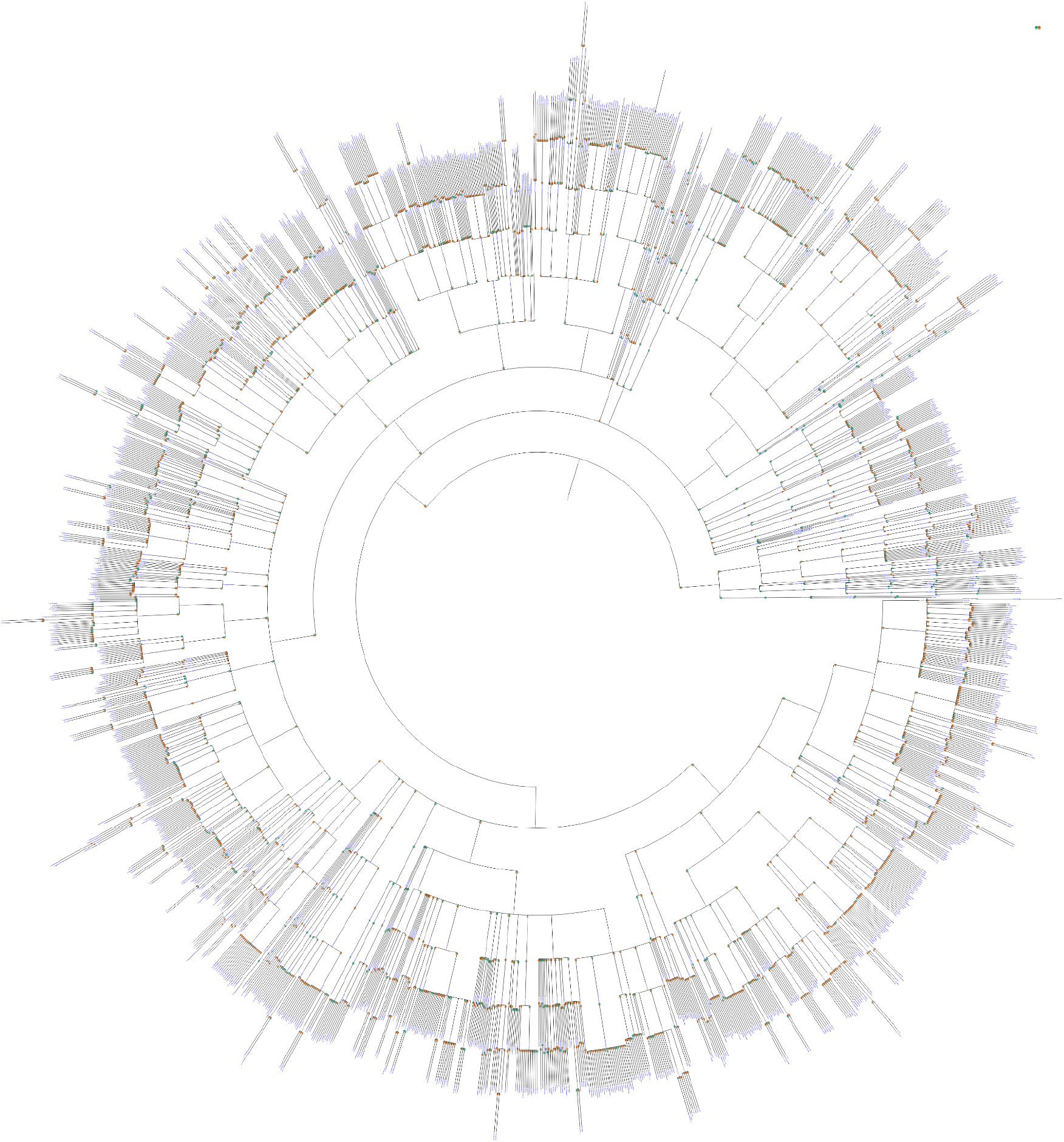
Visualization of the taxonomic profiles of tools with identical UniFrac scores of 4, Taxy_pro vs Metaphyler using TAMPA on the CAMI dataset at the species rank

**Figure S6:**
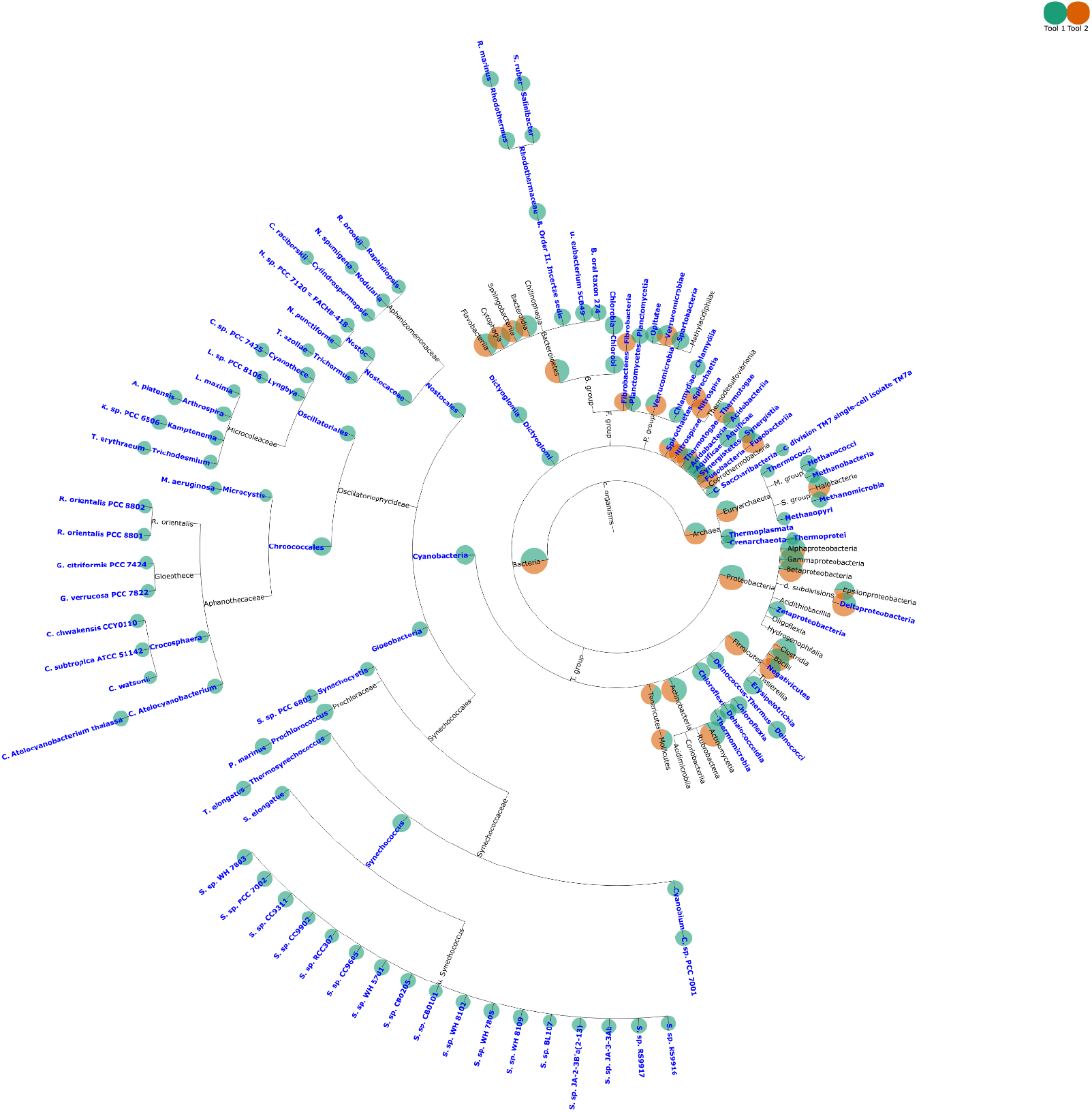
Visualization of the taxonomic profiles of a top performing CAMI tool, Metaphyler vs the ground truth using TAMPA on the CAMI dataset at the class level.

**Figure S7:**
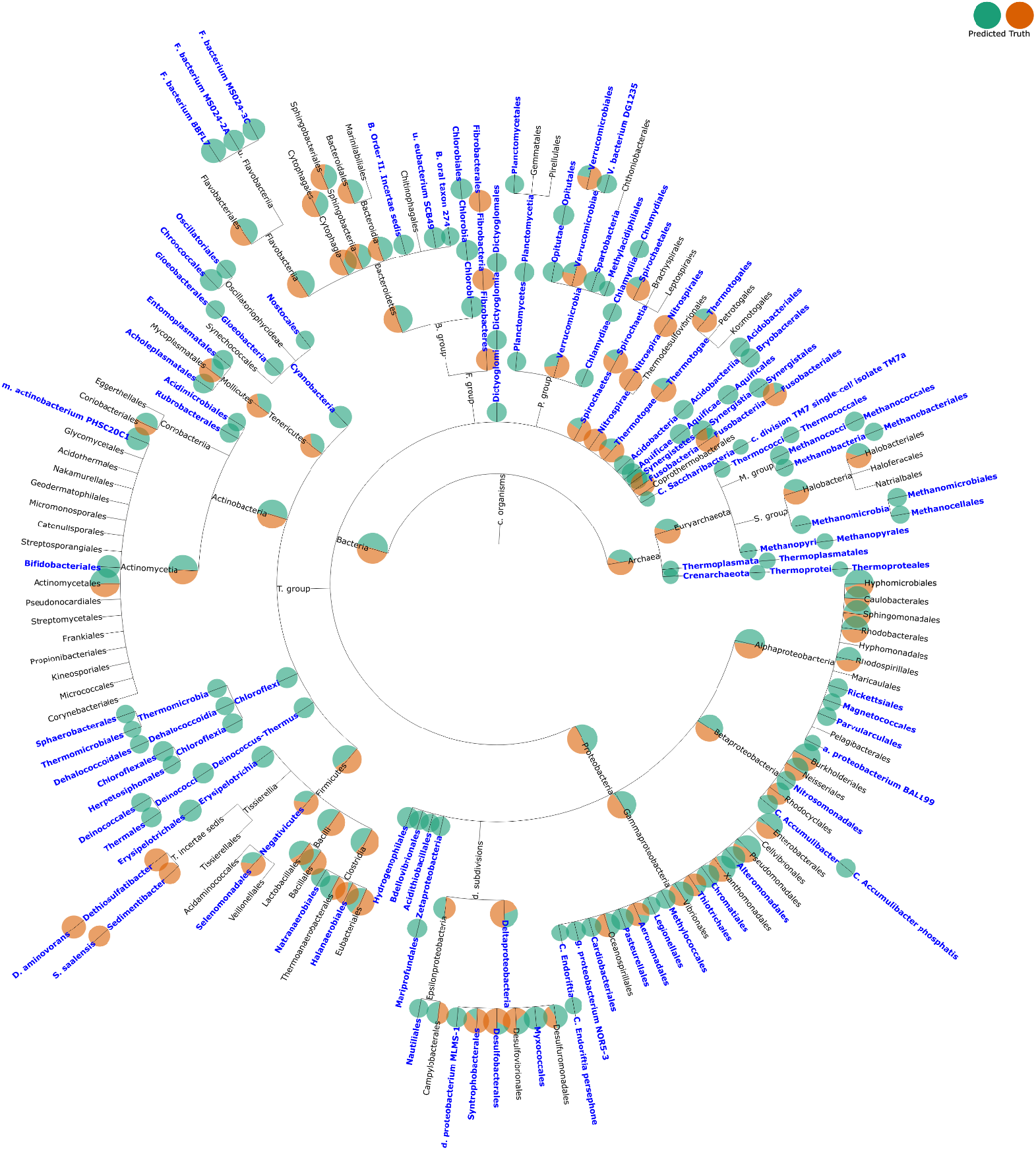
Visualization of the taxonomic profiles of a top performing CAMI tool, Metaphyler vs the ground truth using TAMPA on the CAMI dataset at the order level.

**Figure S8:**
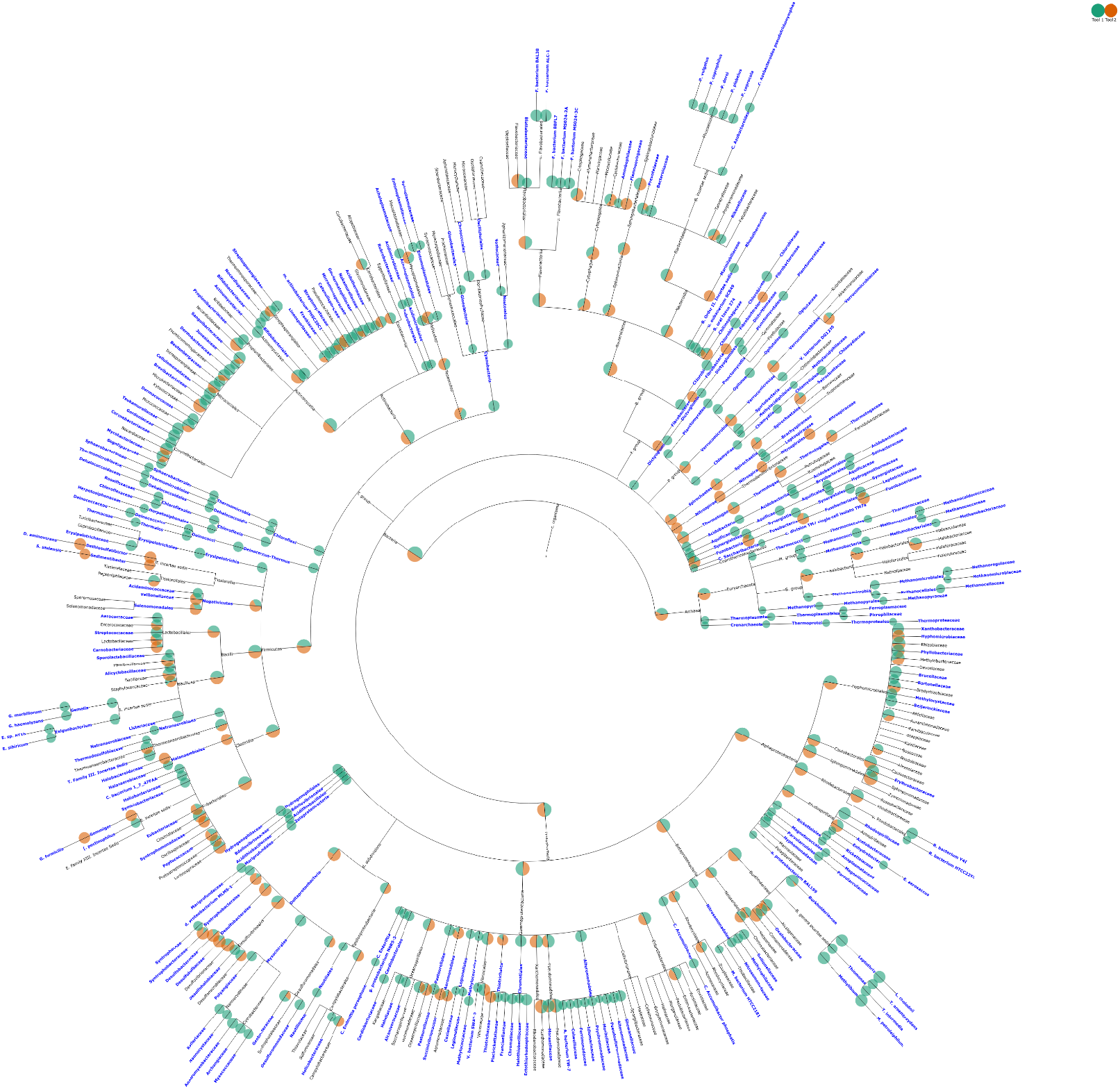
Visualization of the taxonomic profiles of a top performing CAMI tool, Metaphyler vs the ground truth using TAMPA on the CAMI dataset at the family level.

**Figure S9:**
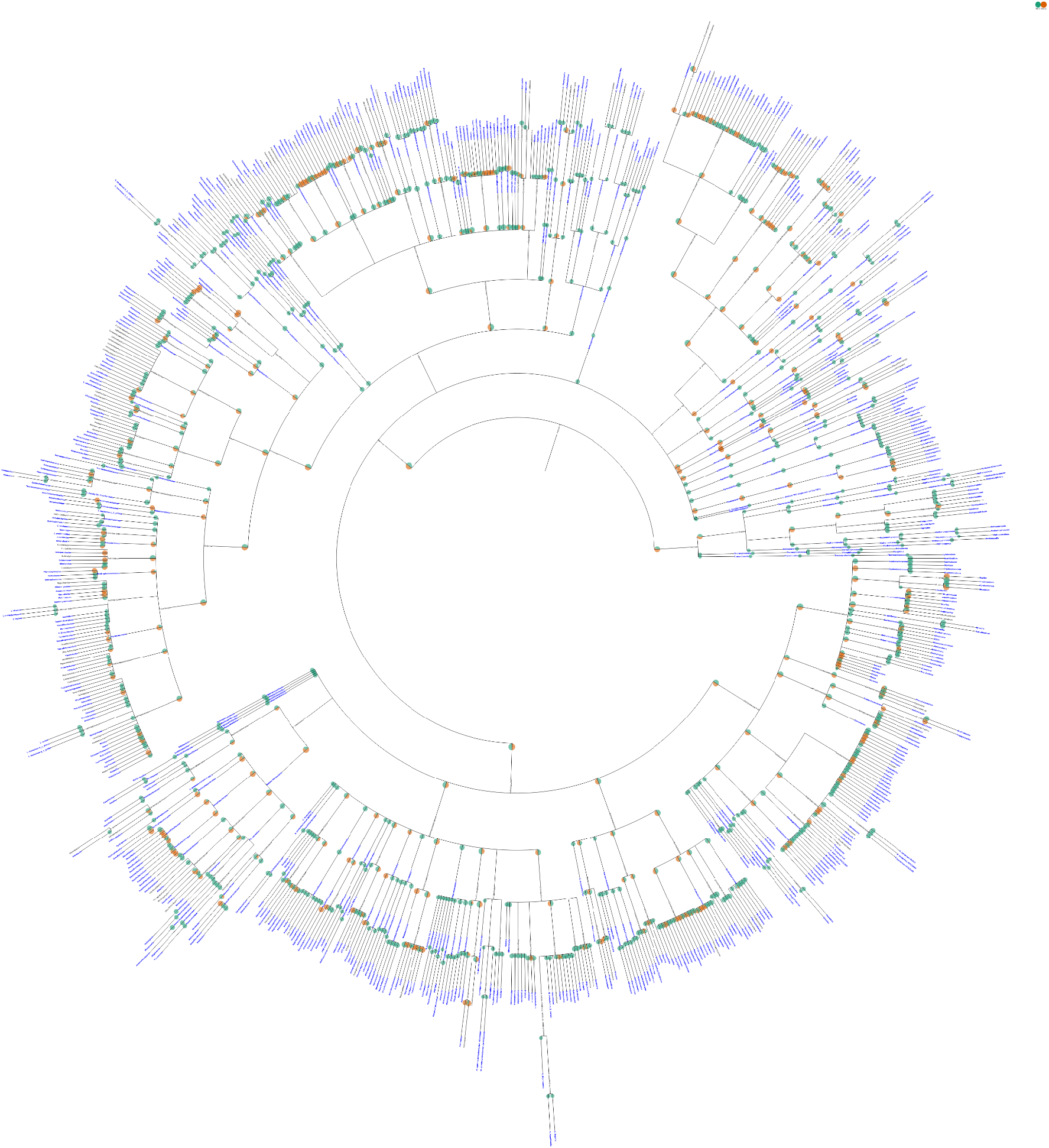
Visualization of the taxonomic profiles of a top performing CAMI tool, Metaphyler vs the ground truth using TAMPA on the CAMI dataset at the genus level.

**Figure S10:**
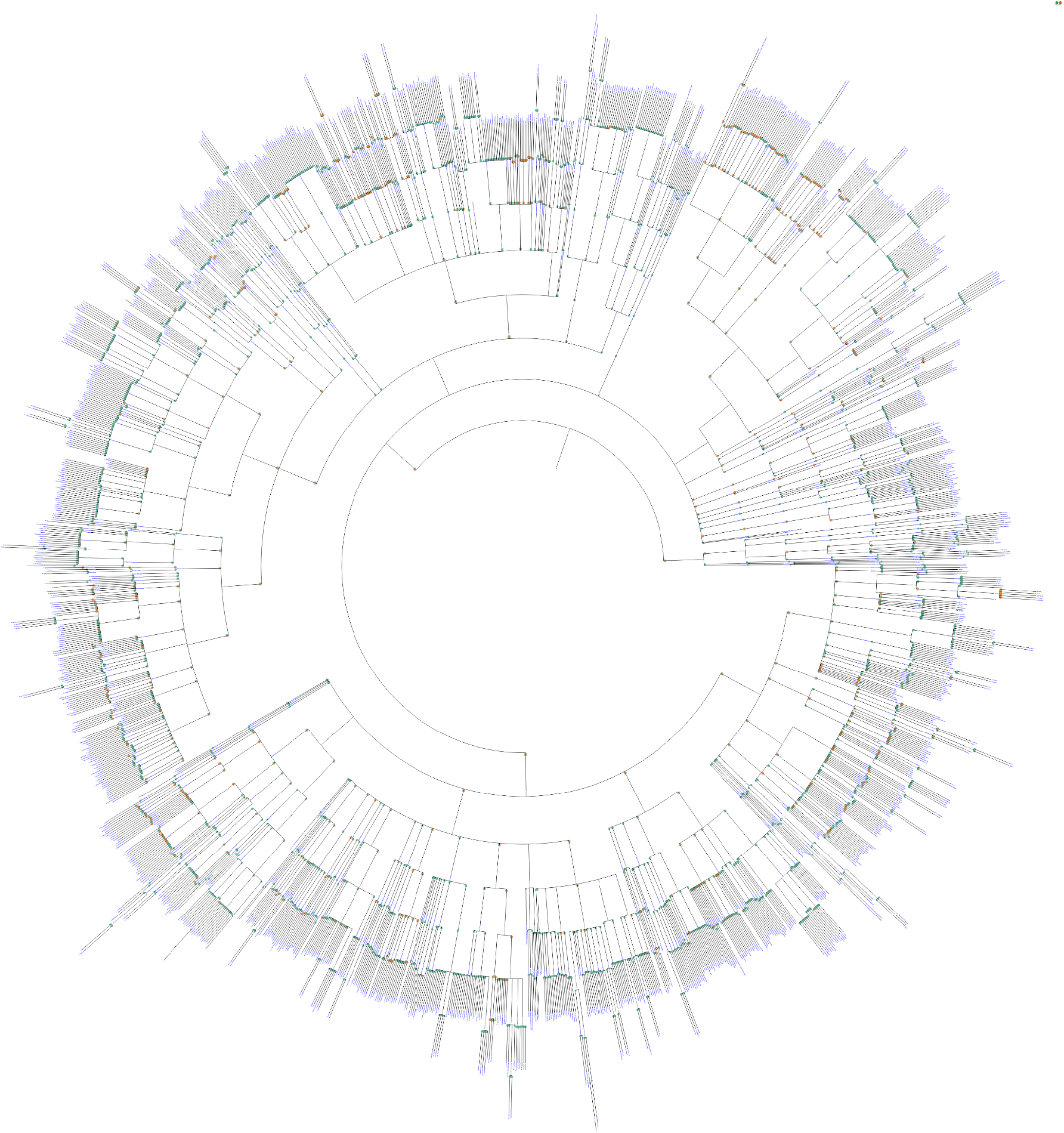
Visualization of the taxonomic profiles of a top performing CAMI tool, Metaphyler vs the ground truth using TAMPA on the CAMI dataset at the species level.

**Figure S11:**
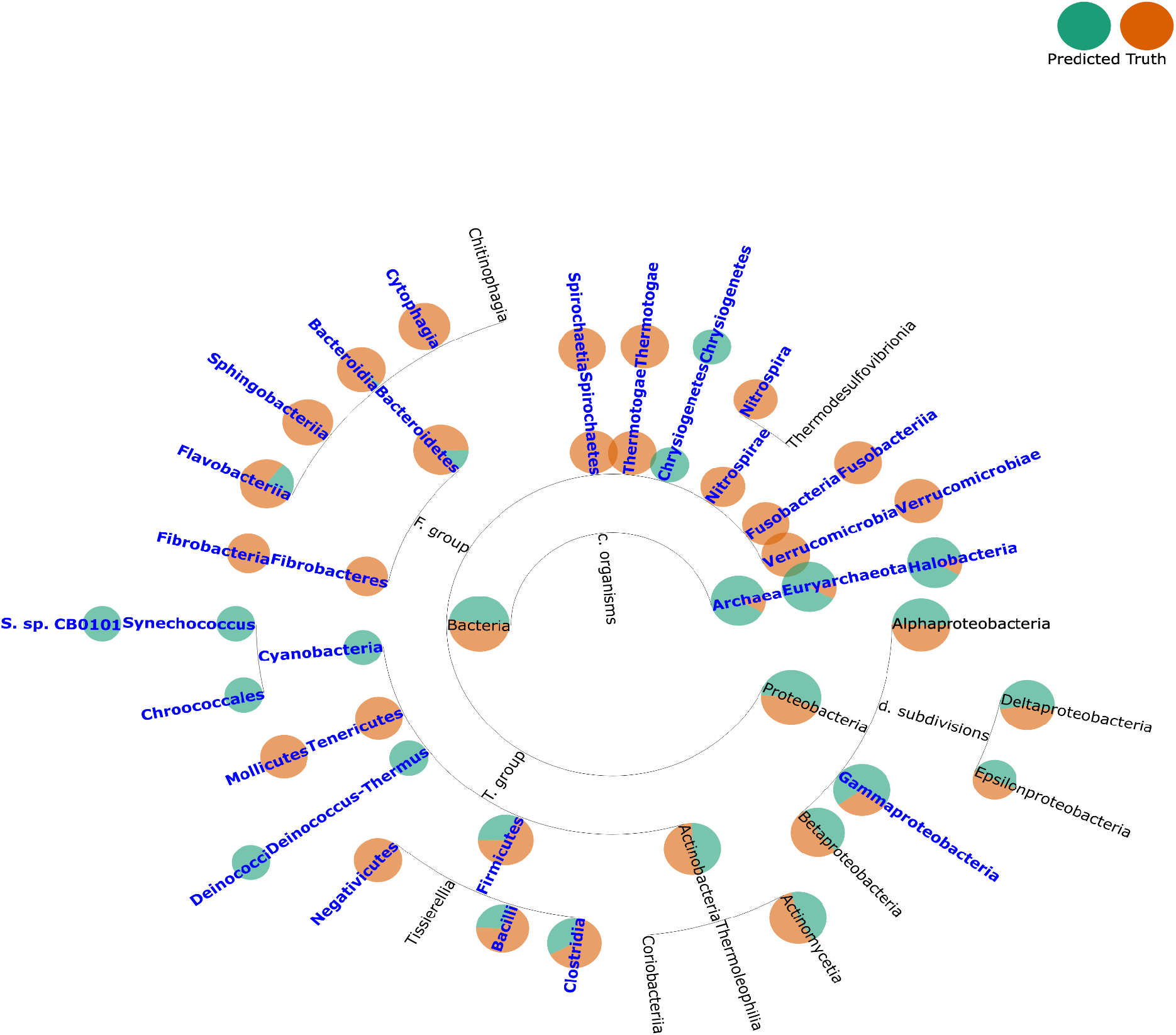
Visualization of the taxonomic profiles of the lowest performing tool, mOTU vs the ground truth using TAMPA on the CAMI dataset at the class level.

**Figure S12:**
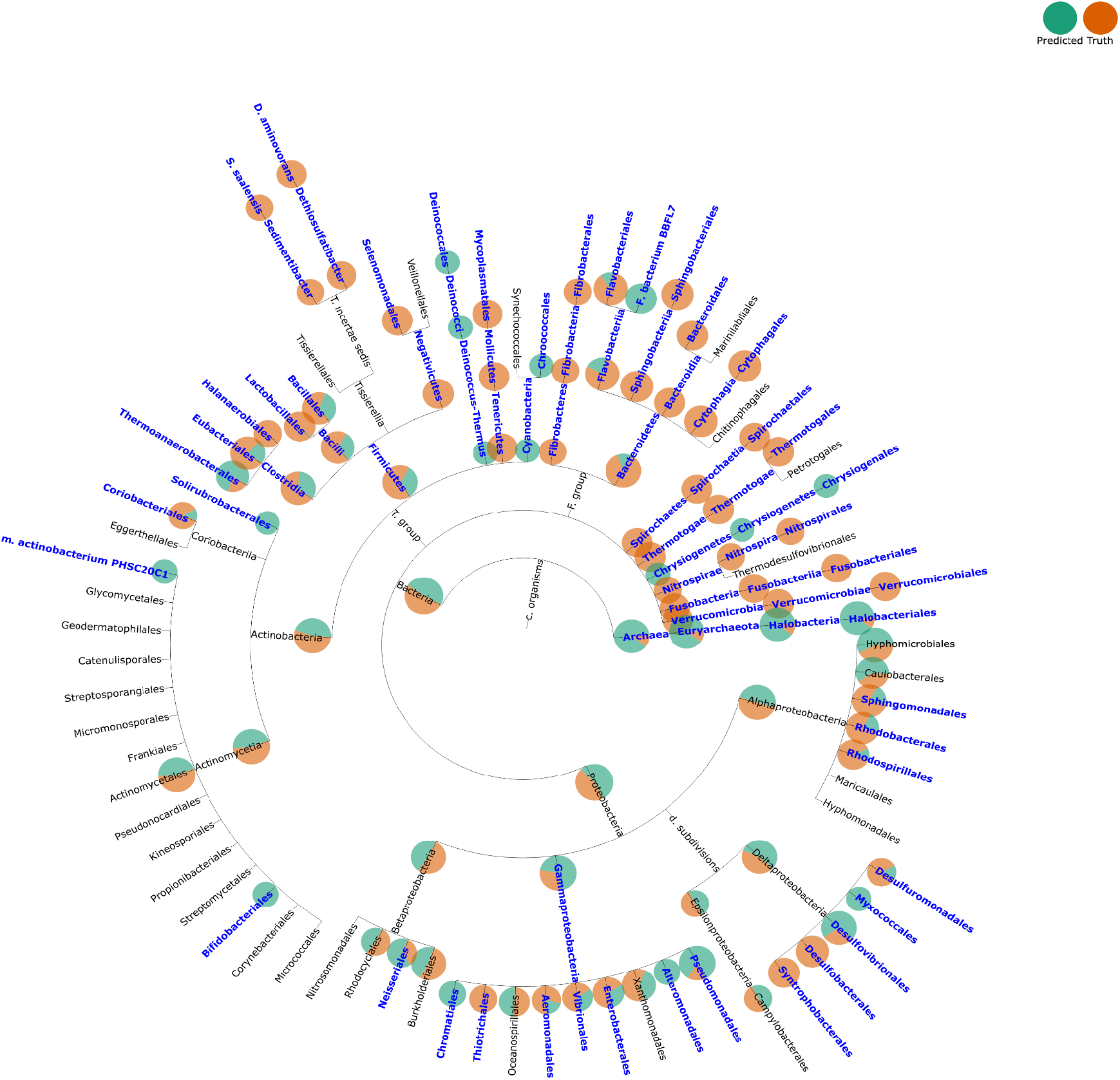
Visualization of the taxonomic profiles of the lowest performing tool, mOTU vs the ground truth using TAMPA on the CAMI dataset at the order level.

**Figure S13:**
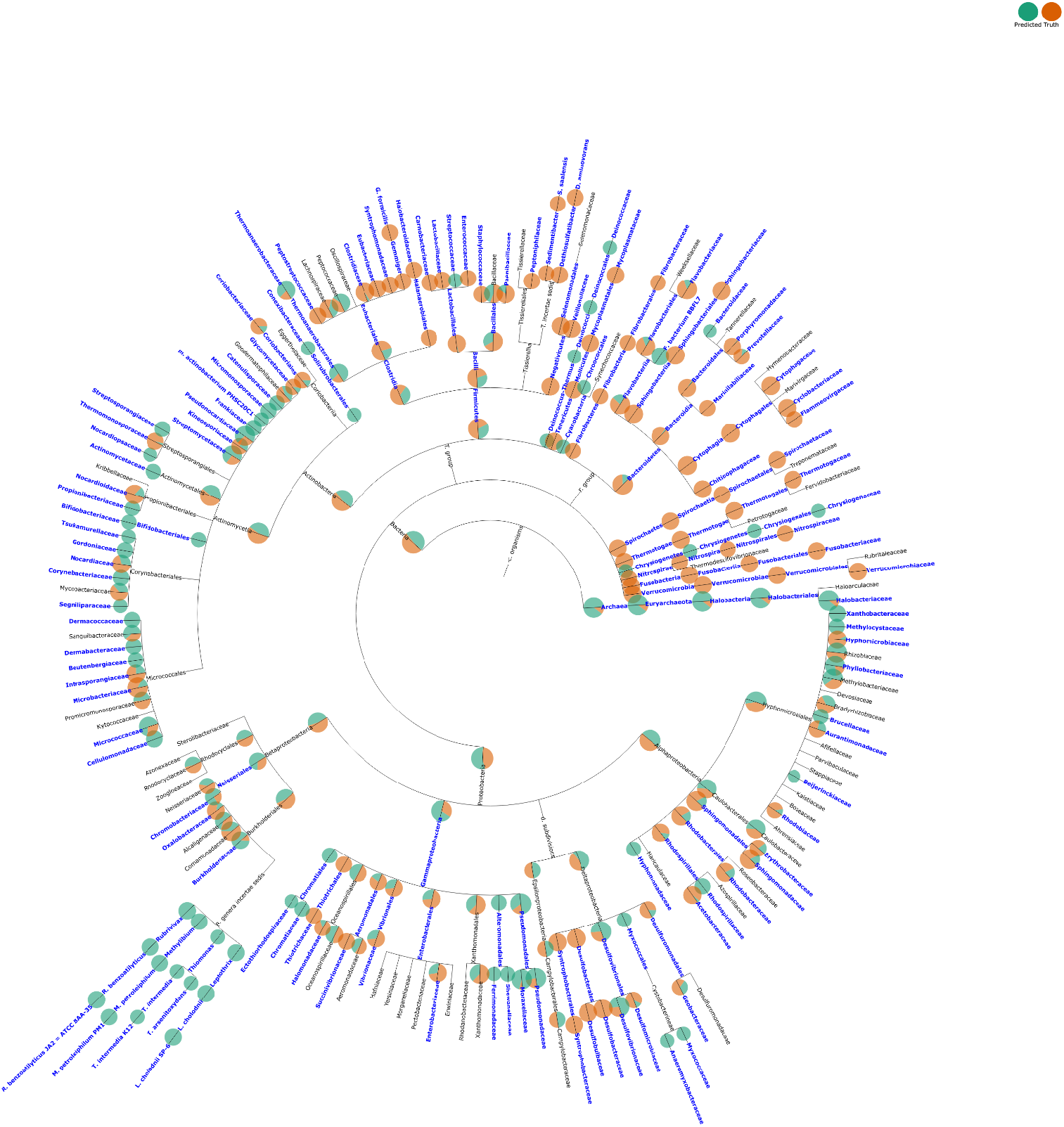
Visualization of the taxonomic profiles of the lowest performing tool, mOTU vs the ground truth using TAMPA on the CAMI dataset at the family level.

**Figure S14:**
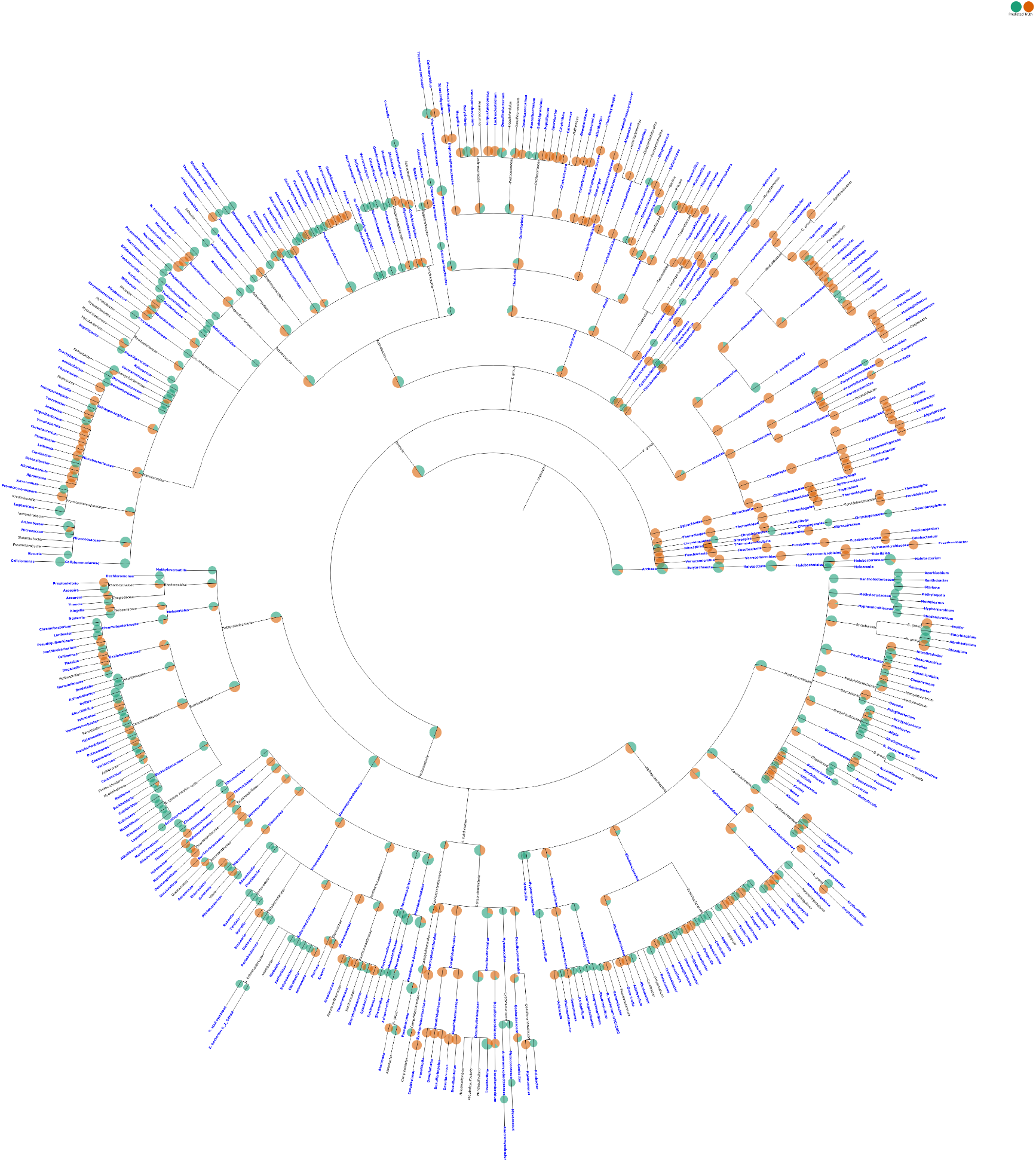
Visualization of the taxonomic profiles of the lowest performing tool, mOTU vs the ground truth using TAMPA on the CAMI dataset at the genus level.

**Figure S15:**
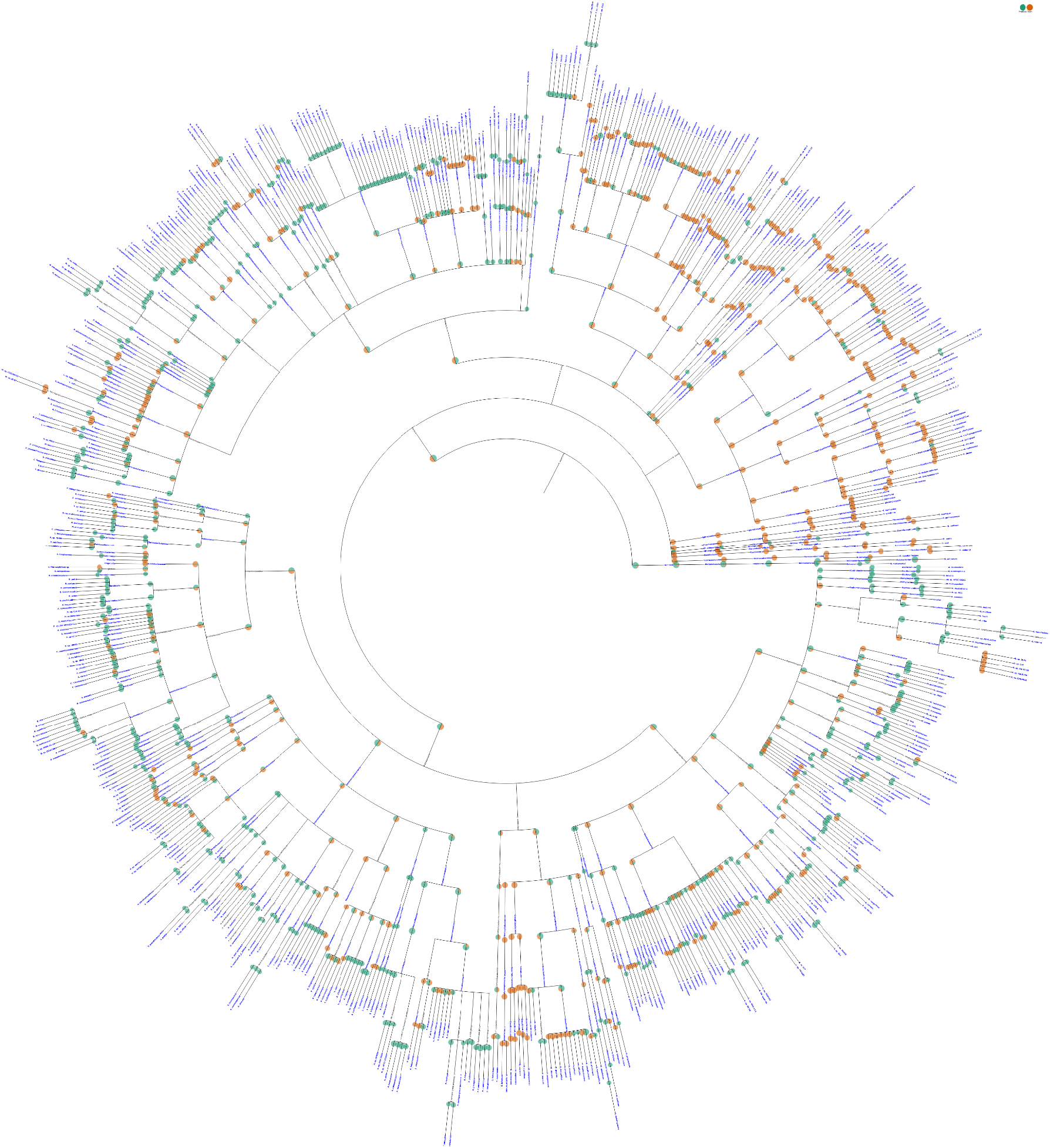
Visualization of the taxonomic profiles of the lowest performing tool, mOTU vs the ground truth using TAMPA on the CAMI dataset at the species level.

**Figure S16:**
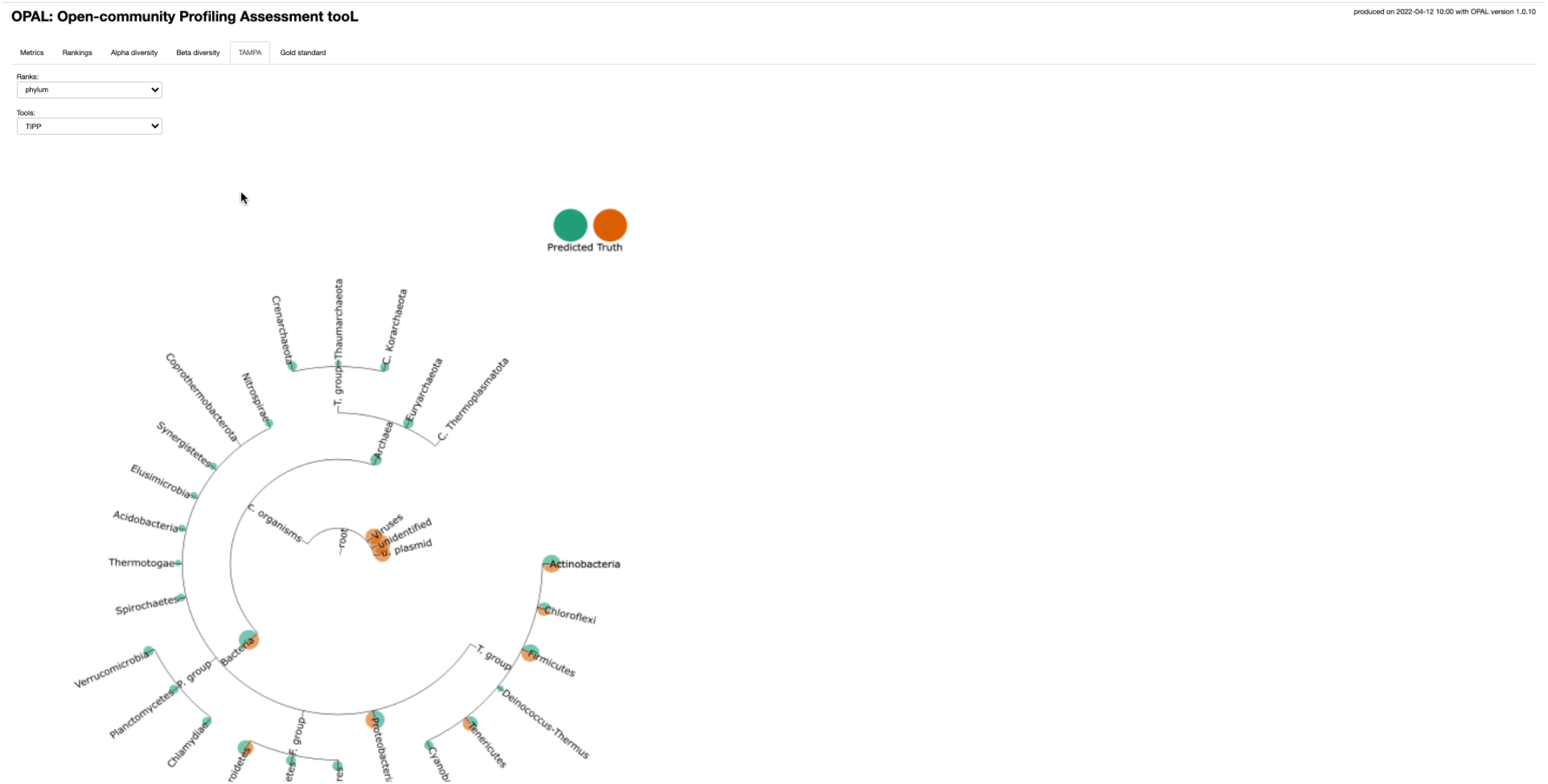
Incorporation of TAMPA into OPAL

